# Differential Recordings of Local Field Potential:A Genuine Tool to Quantify Functional Connectivity

**DOI:** 10.1101/270439

**Authors:** Meyer Gabriel, Caponcy Julien, Paul A. Salin, Comte Jean-Christophe

**Affiliations:** Forgetting and Cortical Dynamics Team, Lyon Neuroscience Research Center (CRNL), University Lyon 1, Lyon, France; Biphotonic Microscopy Team, Lyon Neuroscience Research Center (CRNL), University Lyon 1, Lyon, France; Centre National de la Recherche Scientifique (CNRS), Institut National de la Santé et de la Recherche Medicale (INSERM)

## Abstract

Local field potential (LFP) recording is a very useful electrophysiological method to study brain processes. However, this method is criticized for recording low frequency activity in a large area of extracellular space potentially contaminated by distal activity. Here, we theoretically and experimentally compare ground-referenced (RR) with differential recordings (DR). We analyze electrical activity in the rat cortex with these two methods. Compared with RR, DR reveals the importance of local phasic oscillatory activities and their coherence between cortical areas. Finally, we show that DR provides a more faithful assessment of functional connectivity caused by an increase in the signal to noise ratio, and of the delay in the propagation of information between two cortical structures.

## Introduction

LFP recording of cortical structures constitutes a powerful tool to detect functional signatures of cognitive processes. However, several studies have suggested that recording methods suffer of major caveats due to the recording of activity in distant neural populations [1–4]. Thus, theta oscillations (6-10Hz) during active wake seem to propagate from the hippocampus to the frontal cortical areas [5]. Despite these important studies, LFP recording has revealed important features of cortical organizations [6, 7]. For example, cortical slow wave oscillations of NREM sleep, which constitute a prominent feature of this vigilance state, contribute moderately to coherence between cortical areas [7]. In contrast, weak slow wave oscillations during active wake contribute to a relatively high level of coherence between cortical areas [6, 7]. LFPs are mainly generated by post-synaptic response to pre-synaptic activity of neurons [8–11] and constitutes a natural integrator of action potentials coming from a given cortical region [12–14]. In its usual description, LFP recording appears to be less local than multi-unit activity recordings. Indeed, the usual recording mode of LFP consists in implanting a single electrode in the investigated cortical region and a second one in a supposed neutral site. This simple recording configuration, called monopolar or referential recording (RR) mode, is well adapted to evaluate a global brain state. Unlike single and multi-unit probe, the impedance of the standard electrode used for LFP recording is usually low in order to record neural activity of a larger area. However, this method may detect activities from distant cortical areas located between the recording and the reference electrode [1, 13–19], a phenomenon called *volume conduction*. We propose here to compare monopolar or RR mode to bipolar or differential recording (DR), which consists in setting a pair of electrodes in the same cortical area and measuring the voltage difference between them. The main historical reasons why RR is widely used [7, 20] are: 1) its simplicity because of the low number of wires that needs to be implanted (contributing to the preservation of brain tissue), 2) the number of available channels to connect to the acquisition devices to record the signals, and 3) the method is sufficient to identify global brain states and oscillations in extracellular space. However, to our knowledge, no study has compared both recording methods in freely moving rats in order to define the best suited configuration to record the activity of different brain areas and quantify their interactions, as well as to extract the genuine meaning of the signals recorded in a specific brain region during a behavioral task [1, 7, 20–28]. The present work has been made possible by our recording configuration described in the Methods Section.

Thus, the present paper is organized as follows. First, we present the theoretical *rationale* of the paper. After a description of the experimental conditions, we experimentally show the difference between the two recording modes through spectral analysis and reveal a new communication frequency band between medial prefrontal cortex *PFC* and the dorsal hippocampus area *CA*1. Finally, we numerically show that the assessment of functional connectivity is strongly impacted by the recording mode, indicating why *DR* is much better suited to determine the functional interactions between cortical areas.

## 1 Differential and Referential Recordings

*RR* mode consists in recording the activity of a cortical region by inserting an electrode in the considered (hot spot) area as well as another electrode located in a reference area (ie skull above the cerebellum, cold spot). In contrast, DR consists in setting a pair of electrodes in the same cortical region and to measure the difference of potential between them. In this part, we first analyze theoretically differences existing between the two modes of LFP recordings.

### 1.1 What is volume conduction ?

Volume conduction in brain tissue is a well known phenomenon widely observed in conventional LFP recordings. Volume conduction refers to the process of current flow in a medium. In the brain, the extracellular space contains multiple ionic species.

Even if this biological medium is not really homogeneous, in order to illustrate and simplify our model we consider it as linear, homogeneous and isotropic. Considering a point current source *I* diffusing charges in a sphere of radius *r*, as represented in figure (1), the corresponding density of current 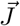 in quasi-static approximation of Mawell’s equations, is given by:

**Fig 1.**
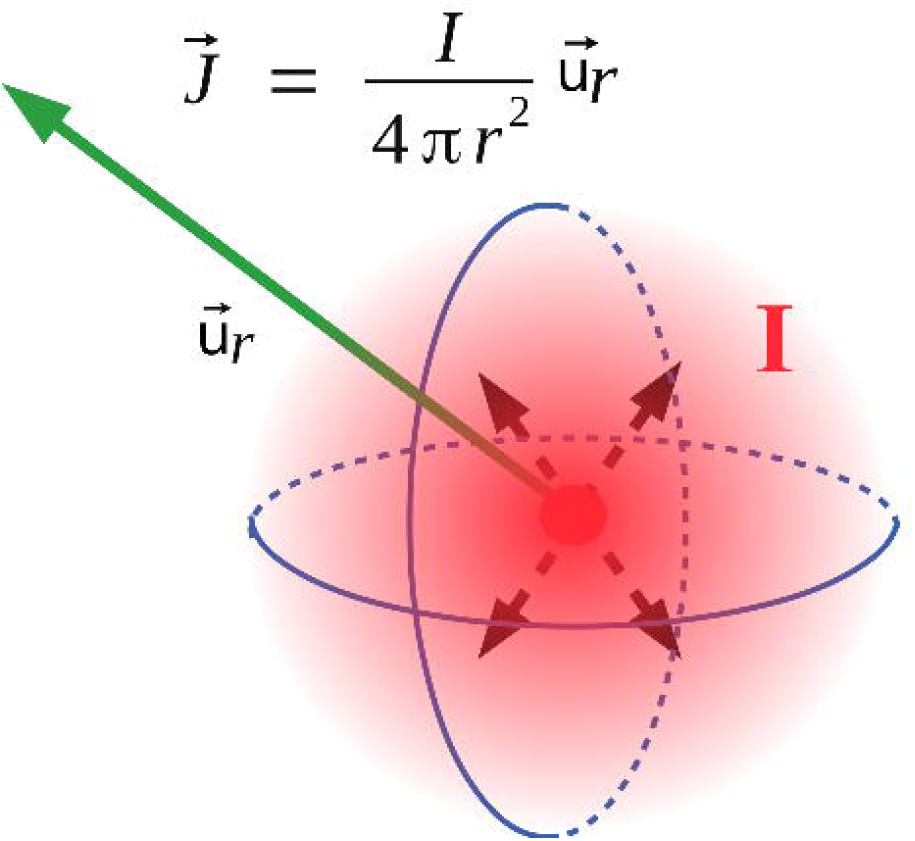
Current source: A current in an homogenous medium yields a current source Iu_r_ density flowing in all directions.The current density writes: 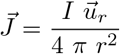 where *r* is the distance to the current source and *I* the current generated at the origin. The Ohm law 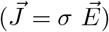 leads to the potential 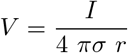 created at any distance *r*.

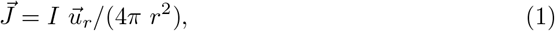

where 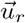 is the radial vector of the current flow direction. Using Ohm law, 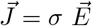 with *σ* being the medium conductivity and 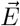 the electric field deriving from the potentiel 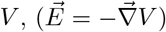, the Potential *V* at a distance *r* is equal to:

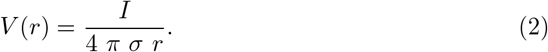

This expression provides the magnitude of the created potential at a distance *r* from a given current source *I*. We observe that this potential decreases nonlinearly with the distance *r*. From this result, we can easily calculate the potential difference between two electrodes *P*_1_ and *P*_2_ separated by a short distance equal to 2ε as represented in figure (2). The potential in *P*_1_ and *P*_2_ is expressed as follows:

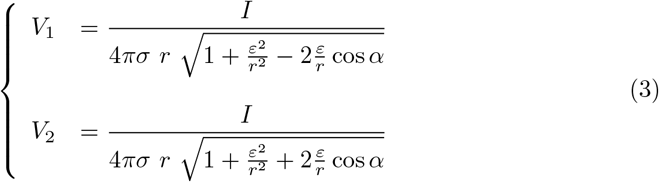

and their difference writes,

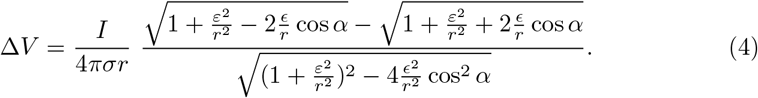

**Fig 2.** a) Distal source: A distal source (blue ellipse) releases a density of current which gives birth to two remote potentials *P*_1_ and *P*_2_ respectivley located at a distance *r* − *δr* and *r* + *δr* belonging to the same brain area (red ellipse). This potential is measured by two electrodes seperated by a distance 2e. b) Local source: A local source (blue ellipse) releases a density of current which gives birth to local potentials *P*_1_ and *P*_2_ respectively located at a distance *r* − *δr* and *r* + *δr* from the source and belonging to the same brain area (red ellipse), where *r* ~ 2*ε*. This potential are measured by two electrodes sperated by a distance 2*ε*.

### 1.2 Case of a distant source

In the particular case *r* >> *ε* (i.e. the distance between an electrode and a source is greater than a few *ε* : in practice *ε* ~ 50-200*μm*), *V*_1_ and *V*_2_ can be rewritten under the form:

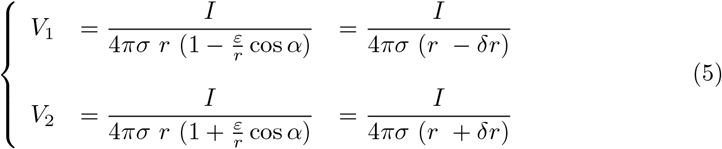

Setting *δr* = *ε* cos *α*, and by neglecting the terms of the second order, the potential difference between the two electrodes writes:

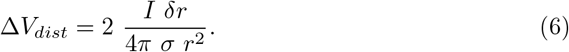

This result shows that adding an electrode in the studied area has the effect of damping the contributions of distant sources by a factor *δr*. Thus, the smaller the distance between the electrodes, the smaller the potential difference. Similarly, the farther a source, stronger is the damping of its intensity. In other words, differential measurement annihilates the contribution of distal sources. We note that, Δ*V_dist_* is maximum for *α* = 0 and minimum for *α* = *π*/2. In other words, the line passing through the two electrodes is the major detection axis.

### 1.3 Case of a local source

Let us consider now the case of a local source contribution, that is, a source close to a a pair of recording electrodes (see fig. 2.b) corresponding to *ε* ≤ *r* < 3 *ε*. Because of the distance between the two electrodes, the minimal distance to a source is *ε*, and when *r* > 3 *ε*, approximations to calculate the potential difference between the two electrodes is similar to the distal source case. As one can observe in figure (2.b), the minimal average distance *r* (electrodes-source) is equal to *ε*, corresponding to a maximal ratio *ε/r* = 1. The ratio *ε/r* < 1/3 yields the ratio *ε*^2^/r^2^ < 1/9 negligible and corresponds to the distant source case. Therefore, to consider the local source case, we approximate *r* to *ε* (*r* ~ *ε*). Under these conditions, the general expression (3) becomes,

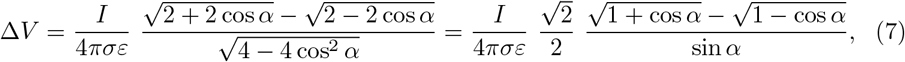

that we note Δ*V_loc_*. Like in the distal case, we note that Δ*V_loc_*(*α* = *π*/2) = 0, while Δ*V_loc_*(*α* = 0) → ∞. In other words, the line passing through the ends of the two electrodes is the major detection axis.

From these results, one can calculate a separation source factor r, or a Common Mode Rejection Ratio (CMRR), by the ratio:

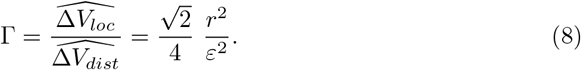

Where 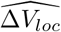 and 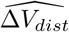, respectively correspond to the maximum of Δ*V_loc_* and Δ*V_dist_*. This factor summarizes that, farther a source, weaker is its contribution. Closer are the two electrodes forming the pair, more visible is the local source. The nonlinearity of this ratio, expressed by the square, indicates that the CMRR rapidly changes with the modification ratio *r/ε*. For instance, for two arbitrary distances *r*_1_ and *r*_2_ equal to 10 e and 100 e respectively, this ratio goes from Γ_1_ = 35 to *Γ*_2_ = 3500, and is σ independant. The present theoretical derivation is true in a ohmic or weakly capacitive extracellular medium approximation. Another formalism should be used [29] to describe a more complex model of the extracellular space, mainly to compare local-local source measurements. Capactive phenomenon can be important at small scale and is neglectable at large scale because of the stochastic distribution of charge in space. Thus, the ohmic approximation of the quasi-static regimes is widely enough to explain the observed differences between RR and DR in the considered frequency range (< 100 *Hz*) and for large distance (>~ 100*μm*) between local and distal sources. More generaly, and intuitively, DR is a mean to remove the common mode visible by a pair of electrode, and consequently allows reveal the difference visible by them.

In both cases, local and distal sources, the maximum voltage detection is obtained for *α* = 0, that is when source is aligned with the two electrodes. This result suggests a better signal detection with four electrodes in a square or at least three electrodes in an equilateral triangle.

Finally, we can summarize all these results in figure (3.a). Figure (3.a) represents the potential measured in *P*_1_ and *P*_2_ *versus* the distance to the source r in normalized units. We note the strong similarity of the potentials when the source is far and their dissimilarity when the source is close. The inset zoom in figure (3.a) shows the strong potential difference between the two electrodes when the source is close to the pair of electrodes. In summary, we have shown that *DR* erases the distal source contribution and constitutes a practical way to solve the volume conduction problem. Even if powerful signal processing methods such as, for instance, partial coherence, may remove signal potential contributions caused by distant neuronal activities [30], an important number of probes would be required to eliminate them as many other cortical areas can potentially generate contaminating signals. Alternatively, in order to avoid volume conduction, it is possible to record the activity of cerebral areas through *DR* using pairs of electrodes in each investigated brain region. In the next part, we assess experimentally the above theoretical predictions and we show the genuine difference between *RR* and *DR* using different tools such as, Fourier analysis, coherence and cross-correlation.

**Fig 3.** a) Example of potential measured in *P*_1_ and *P*_2_ versus distance *r* in normalized units. One notes the strong similitude between *P*_1_ and *P*_2_ when *r* is large in comparison with the distance shift ε of the two electrodes. Also, we oberve a strong amplitude difference between potential P1 and P2 when the current source is close to the electrodes pair (zoom in figure). b) Recording methods and electrodes location during the experiment.

## 2 Experimental Methods and results

In order to verify experimentally our theoretical predictions, we performed LFP recordings in two well known areas of the rat brain, which are the dorsal hippocampus (*CA1*) and the medial prefrontal cortex (*PFC*). The details about the preparation are given in annexe A. Figure (3) shows the recordings configuration in which a pair of electrodes was inserted in each brain region of interest, and a referential electrode was inserted in the skull just above the cerebellum. A calculation of the difference between the two signals coming from the same cerebral structure allows DR. This experimental setup thus enables to compare the two configurations RR and DR modes in the same animal and at the same time. In order to avoid any potential artefacts from the animal movements during wakefulness, we have choosen to focus our attention and analysis on sleep and more specifically on rapid eye movement (REM) sleep (also called paradoxical sleep). REM sleep is characterized by muscle atonia, that can be visualized by a very low power signal on the electromyogram (EMG), and a characteristic cerebral activity visible on the electroencepalogram (EEG) on the form of a low power signal whose spectral energy is mainly located in a narrow band centered around 7 *Hz to* 8 *Hz* (*θ* oscillations). A snippet of such *EEG* epoch is represented in green figure (4.a). Slow wave sleep also called Non rapid eye movements sleep (NREM) is represented in red figure (4). This state was identified by large slow oscillations magnitude accompanied to a low power signal *EMG* but without atonia. Finally, active wake state represented in purple, figure (4.a), presents a low magnitude *EEG* signal close to a gaussian pink colored noise coupled to a strong muscle activity.

**Fig 4.** a) Snippets of typical electroencephalogram (*EEG*) and electromyogram (*EMG*) recordings for the 3 vigilance states, which are wake (Wake→purple), non-rapid eye movement (NREM→red) sleep, and rapid eye movement sleep (REM→green).b) Example of hypnogram showing a temporal vigilance state dynamics.

### 2.1 Spectral analysis.

In order to compare the signal differences between the two recording modes *DR* and *RR*, we performed a spectral analysis by calculating the average power spectrum of the sleep states in *PFC* and *CA*1. Figure (5) shows the power spectra in *RR* mode (blue line) and *DR* mode (red line) in the two investigated brain regions which are CA1 (top), and PFC (bottom), during NREM sleep (left) and REM sleep (right). We should mention that the results presented in this paper were averages obtained from 6 animals, with 195 epochs of NREM sleep and 110 epochs of REM sleep in each animal. Epoch duration has beeen fixed to 15 seconds in order to define a frequency resolution greater than 0.1 *Hz*. The global overview of figure (5) reveals a strong difference between *RR* and *DR* recording modes whatever the brain region and sleep epoch recorded. Beyond the scale factor (~ 10) between the two recording modes, we observe a drastic spectral structure difference. Globally, *DR* spectra present a broader spectral band than *RR*, whatever the brain region and sleep stage. Also, *DR* spectra present a more complex architecture than *RR* spectra. In other words, signals from *DR* and *RR* are qualitatively different even if some parts are similar. Indeed, *RR* is the mix of signals coming from the region of interest as well as signals coming from other asynchronous source regions. Remote asynchronous sources interfere destructively with the local source leading to a rapid decay of the spectrum. *DR* on the other hand, annihilates interfering signals coming from remote sources and then highlights the intrinsic signal of the region of interest as expected by our demonstration in section 1. We can also observe that this fundamental result is state independent. In the next section, we analyze the *CA*1 and *PFC* interplay during REM and NREM sleep in the two recording modes (*DR* and *RR*).

**Fig 5.** Power spectrum of the two simulataneous recording modes *RR* (blue) and *DR* (red). a) and b) respectively corresponds to NREM and REM in *CA*1, while c) and d) respectively corresponds to NREM and REM in *PFC*. In c), arrow shows sleep spindles. b) arrows shows burst activity during *REM* sleep in *CA*1. d) arrow reveals the burst of activity during REM sleep in *PFC*. Note that, burst activity was observed for *DR* in contrast with *RR*.

### 2.2 Coherence and cross-correlation analysis between brain areas.

It is thought that cognitive processes result from information transfer between cortical and subcortical areas [28]. Thus, functional interplay between neuronal populations of different areas remains a major question in neuroscience. Consequently, measurement methods of functionnal connectivity are crucial to test plausible biological hypotheses. We assess functional connectivity, both using *DR* and *RR* mode in the same animal and at the same time to again compare this two modes of recording. We thus calculated the coherence index between *CA*1 and *PFC*. This operation consists in assessing the synchrony or phase locking between two signal sources by expression (9), where *X*(*ν*) and *Y*(*ν*) are respectively the Fourier transforms of two signal sources *x*(*t*) and *y*(*t*). Variable *ν* corresponds to the frequency, while the star sign designates the complex conjugate operator. Coherence index is a statistical tool similar to correlation index but in the frequency domain instead of time. Thus, by this index, we are able to know which spectral component (i.e. frequency) is coherent or phase locked between two cortical areas (cross-spectrum average in the numerator), independently of their magnitude (denominator normalization).

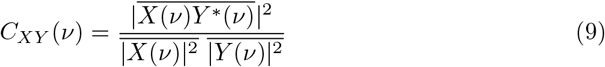

While *RR* and *DR* power spectra of figure (5) share some global common features, figure (6) shows a large difference of coherence between *RR* (blue line) and *DR* (red line), for *NREM* and *REM* sleep. Overall, coherence spectrum appears to be larger using *RR* in comparison with *DR*. The frequency bands in which a peak exists are strongly shifted from one mode (*RR*) to the other (*DR*). For instance, during *NREM* sleep, the frequency peak is located at 1 *Hz* and 3.5 *Hz* respectively, for *DR* and *RR*. Furthermore, during REM sleep, the biggest peak for recording modes *RR* and *DR* are located at 7 *Hz* and 12 *Hz* respectively. These experimental results, confirm that *DR* and *RR* are two different recording modes with their own physical meaning as we demonstrated theoretically in section (1). Unlike *RR*, *DR* gives access to the intrinsic signal of a given cortical area, and therefore to the genuine activity of the investigated neural network. Coherence is a tool that makes sense to assess the functionnal connectivity between two cortical regions. Consequently, it appears that coherence is strongly dependent of the recording mode. It is also important to note that coherence level is not stationary over time. Indeed, as illustrated in figure (7), we oberve that the frequency band 10 *Hz* to 14 *Hz* presents occasional burst of activity in the two recorded cortical structures (*PFC* and *CA*1) at the same time such as the one located at *t* = 20s. However, an oscillation at 7 *Hz* persists all along the REM sleep episode in *CA*1 in the two recording modes. A horizontal projection of this time-frequency diagram provides spectra similar to the figure (5.b) and (5.d), where the average of 7 *Hz* is bigger than the 10 *Hz* to 14 *Hz* in *CA*1, because of the phasic (ie occasional) nature of this 10 − 14*Hz* oscillation. In order to show that *DR* and *RR* modes do not measure the same things, we have also reported the time-frequency of the same period of *CA*1 and *PFC* in *RR* and *DR* mode in figure (7). Even if figure (7.*DR*) and figure (7.*RR*) share some similarities, we can observe that *PFC* recording in *DR* presents no *θ* rythm unlike in *RR* mode (7.*PFC_RR*). We also observe that *DR* shows a power modulation of the low frequency band (< 5 *Hz*) in *CA*1 in contrast to *RR* mode.

**Fig 6.** Coherence index between two brain regions (*CA*1 and *PFC*) during *NREM* a)and *REM* b). Blue lines and red lines respectively correspond to *RR* and *DR*. Arrow in b) show the burst of activity during *REM* sleep. Note that, burst activity was observed for *DR* in contrast with *RR*.

**Fig 7.** Time-frequency representation of a simultaneous *PFC* and *CA*1 recordings in *RR* and *DR* mode during REM sleep, showing an occasional large frequency burst of activity common to the two brain structures located at 20s as well as a persistant oscillation at 7 *Hz*(*θ* rythm) which takes birth in *CA*1. *θ* oscillation is a fundamental REM sleep signature in *CA*1. Colorbar is the normalized scale color of the time-frequency plot. We note that, *θ* rythm is viewable in *PFC* in *RR* mode (*PFC_RR*) in contrast to *DR* mode [*PFC_DR*) showing the volume conduction phenomenon. Occasional burst of activity at 20s is better identified in *DR* (*CA*1_*DR*) mode than *RR* (*CA*1_*RR*) mode.

Finally, it appears that occasional burst is spectrally more extended in *DR* than in *RR* in both areas. For instance, the occasional 10 − 14*Hz* oscillation is simultaneously observed in *CA*1 and *PFC* during REM sleep, but it appears to be bigger with *DR* (figure 7,*PFC_DR* and 7.*CA*1_*DR*). This observation motivates the exploration of the dynamics of the coherence index. Hence, we performed the coherence calculation when a 10 − 14 *Hz* event emerges in one of the two investigated brain structures. In order to perform this analysis, we developed a detection routine allowing to isolate the 10 − 14 *Hz* events. The averages in the coherence plots are thus carried out on the burst events only. Figure (8) shows the coherence factor between *CA*1 and *PFC* during REM sleep. The blue and red traces correspond respectively to *RR* and *DR* mode, while thin and large traces correspond respectively to the triggering area source (*CA*1 or *PFC*). As expected, the choice of the triggering source (*CA*1 or *PFC*) does not change the coherence results whatever the recording mode *RR* or *DR*. The coherence level in *DR* mode is drastically boosted in comparison with the sliding window average method (figure 6) since the level increases from 0.35 to 0.55, while the coherence level in *RR* mode is drastically reduced from 0.6 to 0.45. Moreover, in order to demonstrate that coherence level obtained with *RR* mode is owing to the volume conduction phenomenon, we have calculated the Imaginary Coherence (IC), which ignores the contribution of volume conduction [38]. As shown in figure (8), the two majors peaks in *RR* mode, the one at very low frequency as well as the one located at 7 *Hz* (8.a) are strongly damped when we calculate the IC (8.b), meaning that there is no significant phase shift between cortical areas. Phase shift is due to a propagating phenomenon, while a zero phase shift is due to a conductive phenomenon. The level of these two peaks is reduced to the basal level of the other frequencies, suggesting that IC is altered by volume conduction, since volume conduction is responsible for the real part of the coherence. Another useful measurement to understand how brain areas communicate, is cross-correlation function. This operation is similar to coherence but it is in the temporal domain. It allows to determine the propagation delay between the two investigated brain structures. Propagation direction is determined by the lag sign and the choice of the referential signal (here *PFC*). Figure (9) shows an example of the cross-correlation of two individual burst events (in *DR*) present in *CA*1 and *PFC*. The maximum peak of magnitude 0.55 is 35 ms lagged, that corresponds to a delay of the signal observed in *PFC* in comparison with *CA*1 [39]. In order to compare the ability to measure a delay according to the measurement mode (*RR* versus *DR*), we have performed multiple cross-correlation calculations to construct the lag time probability density function and its corresponding cumulative probability in the two measurement conditions (see fig.9b and c). Figure (9b) indicates a null median lag time for the *RR* mode presenting a fuzzy probability density distribution around zero, while a 35 ms median lag time is observable for *DR* mode presenting a genuine identified peak (fig.9c). This lag time value is comparable to the measure obtained by using single cell recording mode [1, 39] which consists in recording simultaneously one individual neuron in each structure. These kinds of measurements are difficult to perform and allow to probe only one neuron at a time in comparison with LFP which is the superimposition of the effective activity of hundreds of neurons reflecting the entire network activity. LFP consequently avoids performing multiple single cell recording. In summary, *DR* mode is an efficient way to assess the functional connectivity between brain regions and to identify the communication direction, unlike *RR* mode. In order to avoid a “dilution” process through time, occasional communication between brain regions need to be detected, and functional connectivity must be assessed during periods of communication only.

**Fig 8.** a) Coherence between *CA*1 and *PFC* during REM sleep for the two recording modes RR (blue) and *DR* (red). Triggering source: Thin traces correspond to a trigger according to *CA*1, while large traces correspond to a trigger according to *PFC*. b)Imaginary Coherence between *CA*1 – *PFC* in *RR* configuration, showing the decrease of the 7 *Hz* peak as well as the very low frequency peak, because volume conduction is mainly represented by the real part. The 10 *Hz* to 15 *Hz* frequency band stays absent because of the poor signal to noise ratio in *RR* configuration. Inset: vertical zoom of the coherence index.

**Fig 9.** a) Individual event cross-correlation between *CA*1 and *PFC* in *DR* mode, showing a maximum correlation level of 0.55 at a positive lag time of 35 ms between the two regions. This positive lag indicates in our case a delay from the *PFC* in comparison with *CA*1. b) and c) are the probability density functions (blue) and cumulative probabilities (red) of the cross-correlation peak lag. A zoom on the maximum of the probility density function shows in referential mode b) a null median lag time and a fuzzy probility density function, while in differential mode c), the zoom displays a very well indentified peak and median lag time of 35 ms.

Finally, we have performed numerical simulation in order to show the impact of the signal to noise ratio (SNR) on the coherence index measurement. As we suggested above, *RR* mode integrates the contribution of the distal sources weigthed by the distance, while *DR* mode annihilates the contribution of distal sources. Consequently, the SNR is not the same in both configuration. *SNR* is the ratio of the voltage mesured for a local source in comparison with a distal source. In *RR*, the voltages respectively mesured for a local and distal source are

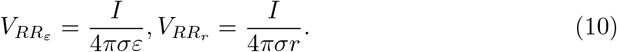

Thus, *SNR* in *RR* mode is the ratio of the wanted signal *V_RR_ε__* and the unwanted noise *V_RR_r__*, that is,

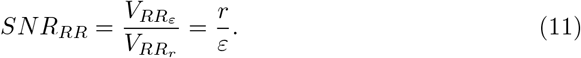

Similarly, in *DR* mode, the *SNR* is the ratio of the local wanted signal *V_DR_ε__* and the distal unwanted noise *V_DR_r__*, and writes:

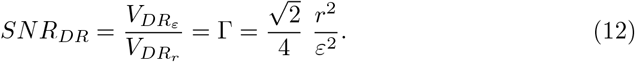

Finally, in order to compare the two SNR corresponding to the *RR* and *DR* mode we define the ratio:

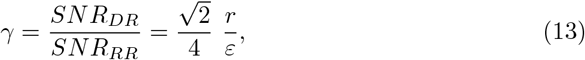

which is greater than one when 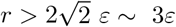 corresponding to the limit between local and distal source as considered in susbsection (1.3). *SNR_DR_* grows faster than *SNR_RR_* proportionally to r and inversely proportional to *ε*. *r* < 3*ε* is the local sphere measurement. Let us consider now the arbitrary choice of a distance equal to two times the radius of the local sphere, that is *r* = 6*ε*. In this case, 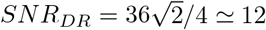, while *SNR_RR_* = 6. This result obtained in one dimension of space leads to a *SNR_DR_* two times greater than *SNR_RR_* for a distance of 150*μm*. Considering the real three dimensions of space, the global SNR writes: *SNR_G_ = SNR_x_ × SNR_y_ × SNR_z_* = 2 × 2 × 2 = 2^3^ = 8. Thus, *DR* mode presents a SNR eight times greater than *RR* mode.

Since *SNR* is strongly different between the two recording modes *RR* and *DR*, we may wonder what could be the impact that *SNR* has on coherence measure. In order to give an answer to this question, we have performed numerical simulations to construct the relation: Coherence Level versus Noise to Signal Ratio (*SNR*^-1^). Figure (10.c) shows the impact of *SNR* on the coherence index level. The two arrows indicates the coherence level obtained when *SNR*^-1^ is equal to 10 and 20 corresponding respectively to figure (10.a) and figure (10.b). As indicated in figure (10.c), coherence level decreases from 0.5 to 0.1 when *SNR*^-1^ increases from 10 to 20. In other words, coherence level decreases 5-fold when noise is simply double. In summary, we have shown that *DR* and *RR* measurements are not similar regarding *SNR*, and that coherence level is strongly dependent of *SNR*. Thus, because of volume conduction, coherence level in *RR* mode is then overestated and masks the true coherence between cortical areas. Indeed, this common signal interferes sometimes constructively and sometimes destructively with the signal of interest, and consequently may underestimate the true coherence between brain areas.

**Fig 10.** Coherence index vs noise to signal ratio (*SNR*^−1^). a) and b): The coherence calculations have been performed between a pure sine wave (green line ) of unit amplitude vs itself added to a gaussian white noise where the magnitude has been chosen to 10 and 20 respectively for a) and b). c) The coherence index decreases rapidely with *SNR*^-1^ according to a hyperbolic secant law (red line). Arrows point out the coherence level corresponding respectively to a *SNR*^-1^ = 10 and 20.

## 3 Discussion

The aim of this paper was to show and explain the differences between the two recording modes, *RR* and *DR*, as well as to examine a way to reduce the impact of volume conduction in the functional connectivity assessment. Consequently, we have theoretically demonstrated that *RR* and *DR* are two recording modes with their own properties. We have shown that *RR* is more suitable to define the global state of the brain because of volume conduction. On the other hand, we have demonstrated that *DR* is able to annihilate the influence of distal sources and is able to probe specific regional activity. Our experimental recordings analysis in the rat show that *DR* makes possible the study of the interplay between brain areas. Indeed, our coherence analysis shows that *CA*1 and *PFC* exhibit a frequency band located between 10 *Hz* and 15 *Hz* which is not present in the *RR* mode. This result highlights the existence of such a frequency band during REM sleep, which is not easily detectable in *RR* mode. This finding constitutes a new functional signature in REM sleep. Futhermore, we have observed that *θ* oscillations in the frequency band (6 *Hz* to 8 *Hz*) present a strong coherence in *RR* mode, whereas in *DR* mode, this band is almost totaly absent, confirming the contamination of the signal recorded by one electrode over a long distance due to volume conduction. This result fully justifies the use of *DR* mode to investigate the question of cortical areas interactions. Also, we have shown through a time-frequency analysis that communication between *CA*1 and *PFC* is sporadic and not continuous as we expected. Based on the fact that this communication is sporadic, we have performed a new estimation of the coherence, revealing an increase of this one in *DR*, unlike in *RR*. Furthermore, we have computed the cross-correlation synchronized on the burst events in the 10 *Hz* to 15 *Hz* band, and we have statistically shown that *PFC* is 30 ms late behind *CA*1 indicating that *CA*1 is the transmitter and *PFC* the receptor. Finally, we have performed numerical simulations in order to illustrate the relationship between coherence level and SNR. This last result explains clearly the reason why *DR* is better suited to evaluate the interaction between cortical areas than *RR*, since *RR* integrates multiple interfering components. Our study plainly desmontrates the real advantage of *DR* in the understanding of brain communication and consequently for studying memory and learning processes. Also, we hope to motivate through this work the use of *DR* to explore cortical communications in future works.

Many electrophysiological recording tools are available to explore functional brain connectivity. Historically, the use of *RR* was justified by two main reasons. The first one is its simplicity because of the low number of wires required to be implanted (that consequently increases brain tissue preservation). The second one is the number of available channels to connect to the acquisition devices to record the signals. Nowadays, *RR* is still used [7, 20] despite the advent of high density linear electrodes [9, 18, 30] allowing to reconstruct the current-source density topology and location (iCSD) [16, 20]. However, when the experimental protocol is more complex because of the number of cortical sites simultaneoulsy explored in a same animal, *RR* should be used with caution. As shown in our study, *RR* and *DR* modes do not provide the same results and consequently these results cannot be interpreted in a similar way.

To our knowledge, no study has compared both recording methods (i.e. *RR* and *DR*) in freely moving rats in order to define the best suited configuration to record cortical areas activity and quantify their interactions, as well as to extract the genuine meaning of signals recorded in a specific cortical region during a behavioral task [7, 20–27]. In this study, we clarify what is possible to assess according to the recording mode. Indeed, as we have shown above, because of volume conduction, *RR* mode integrates the signals coming from everywhere with a weight inversely proportionnal to the distance. Except in the special case where the signal source is close to the electrode and the distal sources are low, the sum of the contribution of distal sources becomes quickly stronger than the local signal. *RR* is relatively interesting to identify global state changes and is widely used for this matter.

However, some studies have used *RR* to quantify functional connectivity between cortical areas [7, 20]. Although, coherence and cross-correlation differences have been observed between vigilance states, our results, as well as others, suggest that *RR* does not measure the true functional connectivity between cortical areas. *DR* mode has a power spectral magnitude 10 to 100 times smaller than *RR* (see fig.5), while *RR* magnitude keep the same order of magnitude whatever the vigilance state, showing that *RR* mode does not allow local recordings because of volume conduction phenomenon. In contrast, we show in the present study that *DR* mode allows local recordings. This is highlighted first by the spectral structure (5.a and 5.b) in *NREM* and *REM* states for which new spectral bands emerge. Also, this result is strengthened by the coherence analysis that draw attention to a new spectral band of interest during *REM* sleep indicating the existence of spindle waves during this sleep stage. Coherence is a fundamental method to explore the relationship between cortical regions in the linear approximation. Even if a cortical structure is forwardly and strongly connected to another one, the second structure receives signal from other cortical areas which induces a response to their stimulation. In this simple linear model, the functional connectivity is only sensible to the *SNR*, that is the power ratio between the signal of interest and the rest, suggesting that true functional connectivity could be systematically underestimated. In other word, functional connectivity obtained in *RR* mode is overestimated because of volume conduction, while *DR* presents a more specific value of functional connectivity. To conclude, we believe that this work will help new studies describe systematically and clearly their recording methods. Our study strongly suggests that, works on correlation, coherence or functional connectivity between brain areas should not be performed in *RR* mode. Finally, we suggest that the most relevant works regarding the interplay between brain areas must be reevaluated if they have been realized using *RR*.

## 4 Annex A

The data used was collected from 6 Dark Agouti male rats (Janvier Labs) aged of 10-15 weeks and weighing between 200-250 grams. After surgery for electrode implanting, they were kept in individual cages in a 12/12h (9am-9pm) light/dark cycle with ad libitum access to food and water. One week after surgery, the rats were introduced in their recording chamber and plugged for recording. The recording chamber consisted of a 60×60×60cm faradized box with removable container for the litter, so that the rats could be changed daily at 10 am without being unplugged. While in the recording chambers, the animals were exposed to a white noise of 70dB and were also provided with food and water ad libitum. The temperature of the chambers was regulated at 23° C. Once the responses were stabilized, and after at least two days of habituation, baseline recordings, which we used for our analysis, took place during at least 24 hours. The animal care and treatment procedures were in accordance with the regulations of the local (Lyon 1 University CE2A-UCBL 55) and European (2010/63/EU) ethics committee (Approval Number : DR2016-29) for the use of experimental animals. Every effort was made to minimize the number of animals used and any pain and discomfort occurring during surgical or behavioral procedures. The recording pair of electrodes consisted of two twisted tungsten wires (25*μm* in diameter - California Fine Wire, U.S.A.) de-insulated at the tip along approximately 50*μm*. Muscle activity (EMG) in the neck was recorded with a pair of electrodes that were made by gold plating a small and round solder ball at the de-insulated and hooked tip of a conventional small electric wire. In addition, two 100*μm* diameter stainless steel electrodes were implanted for electrical stimulation in the brain, in order to study the synaptic transmission between the hippocampus and the medial prefontal cortex and between the CA3 and CA1 areas of the hippocampus. All these electrodes, along with reference screws, were connected to a custom-made 16 channels analog preamplifier by the EIB-27 connector (Neuralynx U.S.A.). The signals were then conveyed via a rotating connector (Plastics One, U.S.A.) to a 16 channel amplifier (AM-Systems, U.S.A.) within which this signal was amplified with a gain of 1000. Signals from the different electrodes were then acquired and digitized at 5k*Hz* by a custom Matlab software (The MathWorks, U.S.A.) driving a NI-6343 acquisition board (National Instruments, U.S.A.) before being stored on a computer.

## Acknowledgments

We would like to thanks G. Malleret for its help regarding the manuscript correction.

